# The genomic basis of colour pattern polymorphism in the harlequin ladybird

**DOI:** 10.1101/345942

**Authors:** Mathieu Gautier, Junichi Yamaguchi, Julien Foucaud, Anne Loiseau, Aurélien Ausset, Benoit Facon, Bernhard Gschloessl, Jacques Lagnel, Etienne Loire, Hugues Parrinello, Dany Severac, Celine Lopez-Roques, Cecile Donnadieu, Maxime Manno, Helene Berges, Karim Gharbi, Lori Lawson-Handley, Lian-Sheng Zang, Heiko Vogel, Arnaud Estoup, Benjamin Prud’homme

## Abstract

Many animal species are comprised of discrete phenotypic forms. Understanding the genetic mechanisms generating and maintaining such phenotypic variation within species is essential to comprehending morphological diversity. A common and conspicuous example of discrete phenotypic variation in natural populations of insects is the occurrence of different colour patterns, which has motivated a rich body of ecological and genetic research^1–6^. The occurrence of dark, i.e. melanic, forms, displaying discrete colour patterns, is found across multiple taxa, but the underlying genomic basis remains poorly characterized. In numerous ladybird species (Coccinellidae), the spatial arrangement of black and orange patches on adult elytra varies wildly within species, forming strikingly different complex colour patterns^7,8^. In the harlequin ladybird *Harmonia axyridis*, more than 200 distinct colour forms have been described, which classic genetic studies suggest result from allelic variation at a single, unknown, locus^9,10^. Here, we combined whole-genome sequencing, population genomics, gene expression and functional analyses, to establish that the gene *pannier* controls melanic pattern polymorphism in *H. axyridis*. We show that *pannier*, which encodes an evolutionary conserved transcription factor, is necessary for the formation of melanic elements on the elytra. Allelic variation in *pannier* leads to protein expression in distinct domains on the elytra, and thus determines the distinct colour patterns in *H. axyridis*. Recombination between *pannier* alleles may be reduced by a highly divergent sequence of ca. 170 kb in the *cis*-regulatory regions of *pannier* with a 50 kb inversion between colour forms. This likely helps maintaining the distinct alleles found in natural populations. Thus we propose that highly variable discrete colour forms can arise in natural populations through *cis*-regulatory allelic variation of a single gene.

---

Ladybird species have long been studied by geneticists and evolutionary biologists to investigate the origin and maintenance of discrete colour pattern forms in natural populations^7^. In particular, the harlequin ladybird *H. axyridis* is an emblematic species of elytral colour pattern polymorphism, with more than 200 colour pattern forms described from different localities^11,12^. However, four forms dominate natural populations with high frequencies (Fig. 1a)^13^: three distinct melanic forms harbouring different patterns (from darkest to lightest, form *conspicua*, f. *spectabilis*, and f. *axyridis*, and hereafter called Black-2Spots, Black-4Spots and Black-nSpots, respectively), and a non-melanic form (f. *succinea*, called Red-nSpots). The striking array of colour patterns documented in *H. axyridis* in the wild has been attributed to a combination of allelic diversity, interactions between allelic forms, and plastic response to environmental factors^12,13^. Genetic crosses have demonstrated that the majority of *H. axyridis* melanic forms result from variation of multiple alleles segregating at a single, uncharacterised, autosomal locus^9,10^, hereafter referred to as the colour pattern locus.

**Figure 1.**
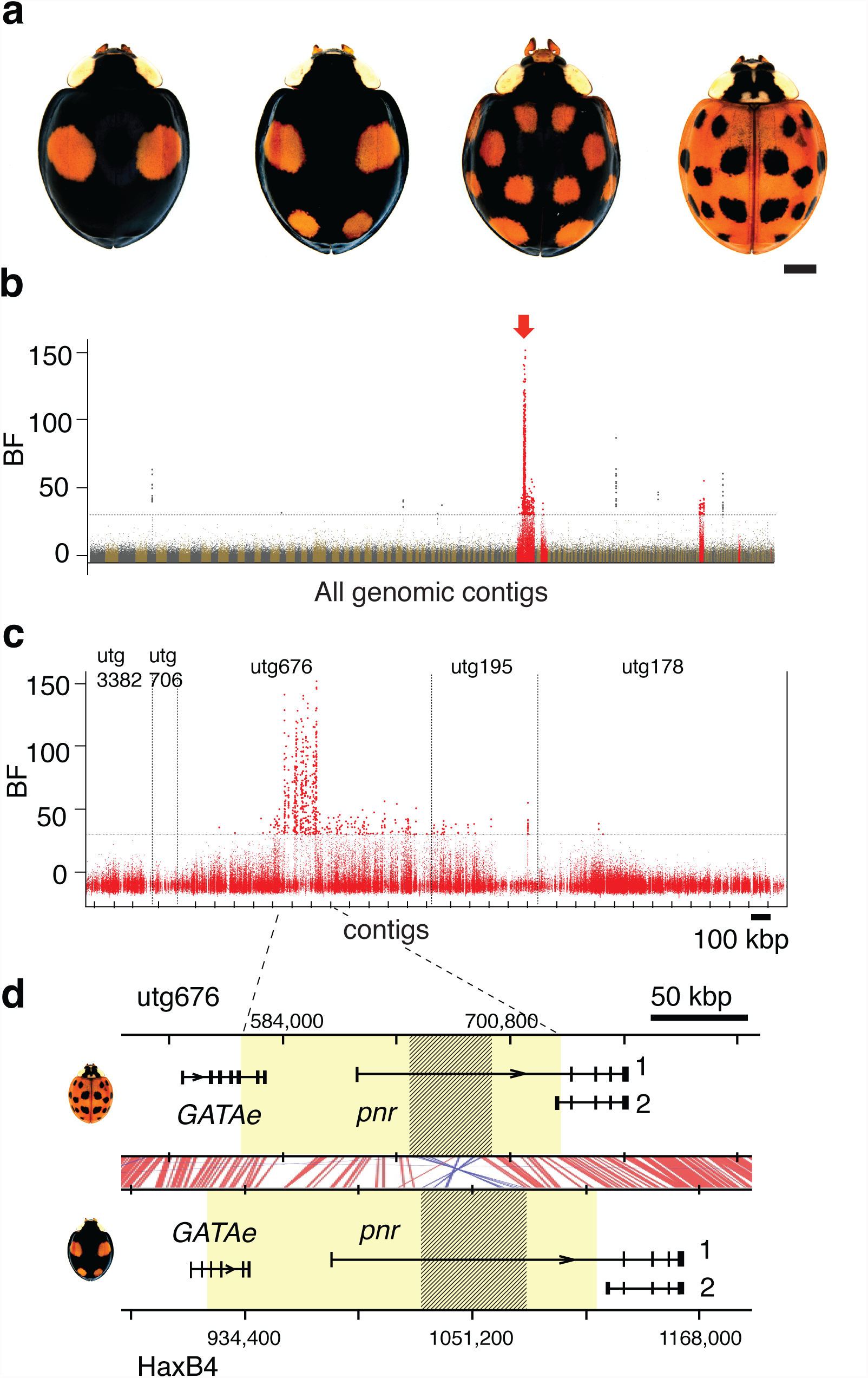
Genome-wide association study identifies the main colour pattern locus in *H. axyridis*. **a**, The four most frequent colour pattern forms of *H. axyridis*. From left to right: the form Black-2Spots (f. *conspicua*), Black-4Spots (f. *spectabilis*), Black-nSpots (f. *axyridis*), and Red-nSpots (f. *succinea*). **b**, Manhattan plots of genome-wide association for the proportion of Red-nSpots individuals in 14 DNA pooled samples of wild H. axyridis populations, with Bayes Factor (BF) for individual SNPs. The horizontal dashed line indicates the 30 db threshold. SNPs above this threshold are highlighted and those assigned to contig utg676 (red arrow, containing the colour pattern locus) in the HaxR assembly and four neighbouring contigs are shown in red. Contigs are ordered by length. **c**, Same as Fig 1b with a focus on SNPs belonging to the five neighboring contigs including and surrounding the colour pattern locus of the HaxR assembly (in red in Fig 1b.). The relative ordering of these contigs was derived from the *de novo* sequencing of the Black-4Spots allele extended region (Methods). **d**, The gene content at the identified colour pattern locus. Fifty-six SNPs with the strongest association signal delimits a candidate colour pattern locus region of ca. 170 kb. (yellow boxes) that extends from the first coding exon of *pannier* to the 5’ upstream gene *GATAe*. Red and blue lines show conserved sequence blocks in forward or reverse direction, respectively, detected by24. The first intron of *pannier* contains the footprint of a ca 50 kb inversion (shaded boxes).

To identify this colour pattern locus, and the mechanisms underlying discrete colour pattern variation, we used a population genomics approach, taking advantage of the cooccurrence of multiple colour pattern forms in natural populations. To that end, we first performed a *de novo* genome assembly of the *H. axyridis* Red-nSpots form (*HaxR*) using long reads produced by a MinION sequencer (Oxford Nanopore) (see Extended Data Table 1). Then, to fine map the colour pattern locus on this assembly, DNA from 14 pools of individuals (from n=40 to n=100 individuals per pool) representative of the world-wide genetic diversity and the four main colour pattern forms of *H. axyridis* was sequenced on a HiSeq 2500 (Illumina, Inc.) (Extended Data Table 2). Our aim was to characterize genetic variation associated with phenotypic differences across pool samples, using the proportion of individuals of a given colour form in each pool as a covariate. To do so, we called 18,425,210 autosomal SNPs, and we performed a population-based genome-wide association study accounting for the covariance of allele frequencies across pools^14^. Given the dominance hierarchy of colour morph alleles with Black-2Spots > Black-4Spots > Black-nSpots > Red-nSpots^9,12,13^, we first performed the association study using the proportion of Red-nSpots individuals, carrying two copies of the most recessive allele, in each DNA pool as a covariate. We found 710 SNPs strongly associated with the proportion of the Red-nSpots form (Bayes factor > 30 db), the vast majority (86%) of which are located within a single 1.3 Mb contig, utg676 (Fig 1b). The 56 SNPs with the strongest association signals (Bayes factor > 100 db) delineate a ca. 170 kb region on *HaxR*, representing the strongest candidate region for the colour pattern locus. Importantly, additional genome-wide association studies using the proportions of Black-4Spots, Black-2Spots or Black-nSpots individuals in the pools as covariates pointed to the same region (Extended Data Fig. 1 and 2), although these analyses were less powerful.

The candidate colour pattern locus extends from the first coding exon of the ortholog of the *Drosophila* gene *pannier*, including its first intron and first non-coding exon, to the end of the neighbouring 5’ gene, the *GATAe* ortholog (Fig 1d). To test a possible role of *pannier* or *GATAe* in adult colour pattern formation, we used RNA interference^15^. Because adult pigmentation patterns are specified during pupal development in insects, we injected larvae of the different *H. axyridis* forms, just before pupation, with double-stranded RNA (dsRNA) targeting the coding sequences of *pannier* or *GATAe*. We also targeted *eGFP* as a negative control. Targeting *GATAe* or *eGFP* had no effect on pigmentation, both in the Red-nSpots and the Black-4Spots forms, suggesting that *GATAe* does not play any role in elytral pigmentation (Extended Data Fig. 3). In contrast, knocking-down *pannier* dramatically reduced the formation of black pigment in all different forms, resulting in adults with almost homogeneous red/orange elytra (Fig. 2). Dark pigment formation is not only strongly reduced in the elytra but also in the head and the rest of the body (Extended Data Fig. 4). Using a different, non-overlapping dsRNA fragment of *pannier* produced similar results, ruling out RNAi off-target effects (Extended Data Fig. 5). These results show that *pannier* is necessary for the formation of black pigment in *H. axyridis* adults. Furthermore, combined with our genome-wide association study, our data indicate that *pannier* is the main gene responsible for colour pattern polymorphism in *H. axyridis,* and that different *pannier* alleles determine the colour pattern in the different forms.

**Figure 2.**
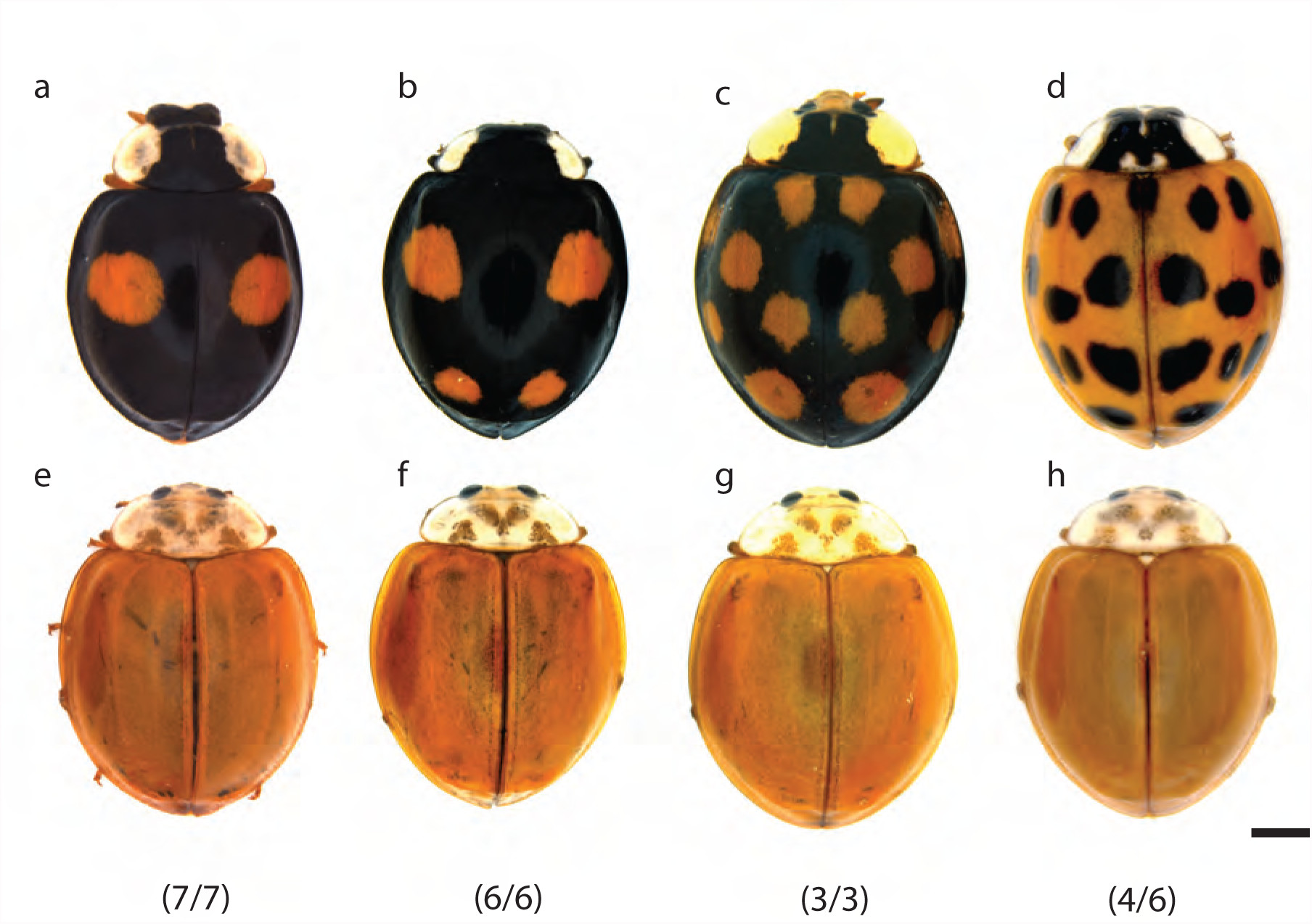
pannier is necessary for black pigment production in *H. axyridis*. Wild type colour pattern forms (upper panels) and representative phenotypes for each form when knocking down pannier by larval RNAi (lower panels). **a**, **e**, Black-2Spots; **b**, **f**, Black-4Spots; **c**, **g**, Black-nSpots; **d**, **h**, Red-nSpots. Numbers indicate the fraction of eclosed adults with the representative phenotype for each form. Scale bar, 1mm.

To understand how *pannier* contributes to the formation of different colour patterns we compared its coding sequences between the Red-nSpots and Black-4Spots forms. We did not find any non-synonymous mutation (Extended Data Fig. 5), thus ruling out changes in Pannier protein composition. We next hypothesised that *pannier* might have evolved divergent expression patterns during the development of the elytra resulting in different colour pattern forms. We therefore compared *pannier* expression level by RT-qPCR in late developing pupal elytra between Red-nSpots and Black-4Spots forms. We found that *pannier* is expressed at a higher level in the elytra of the Black-4Spots form compared to the Red-nSpots form (Fig. 3a). In order to determine how this difference reflected in Pannier spatial expression pattern, we compared the relative *pannier* expression levels in different parts of a Black-4Spots elytron. We found that *pannier* is expressed at a higher level in a presumptive black area, in the middle of the elytron, compared to the red presumptive areas (Fig. 3b). In order to map these differences onto spatial expression patterns, we stained late pupal elytra with an antibody raised against *H. axyridis* Pannier. We found that Pannier spatial distribution on the elytra is different between colour pattern forms (Fig. 3c-f, Extended Data Fig. 6). Strikingly, in all forms, areas with the strongest Pannier expression levels prefigure the adult elytral pattern of melanic elements. This tight spatial correlation, coupled with our genomic association study and the essential role of *pannier* in governing melanic patterns, provide strong evidence that *cis*-regulatory changes at the *pannier* locus drive divergent *pannier* expression patterns and, in turn, the polymorphic melanic patterns of *H. axyridis*.

**Figure 3.**
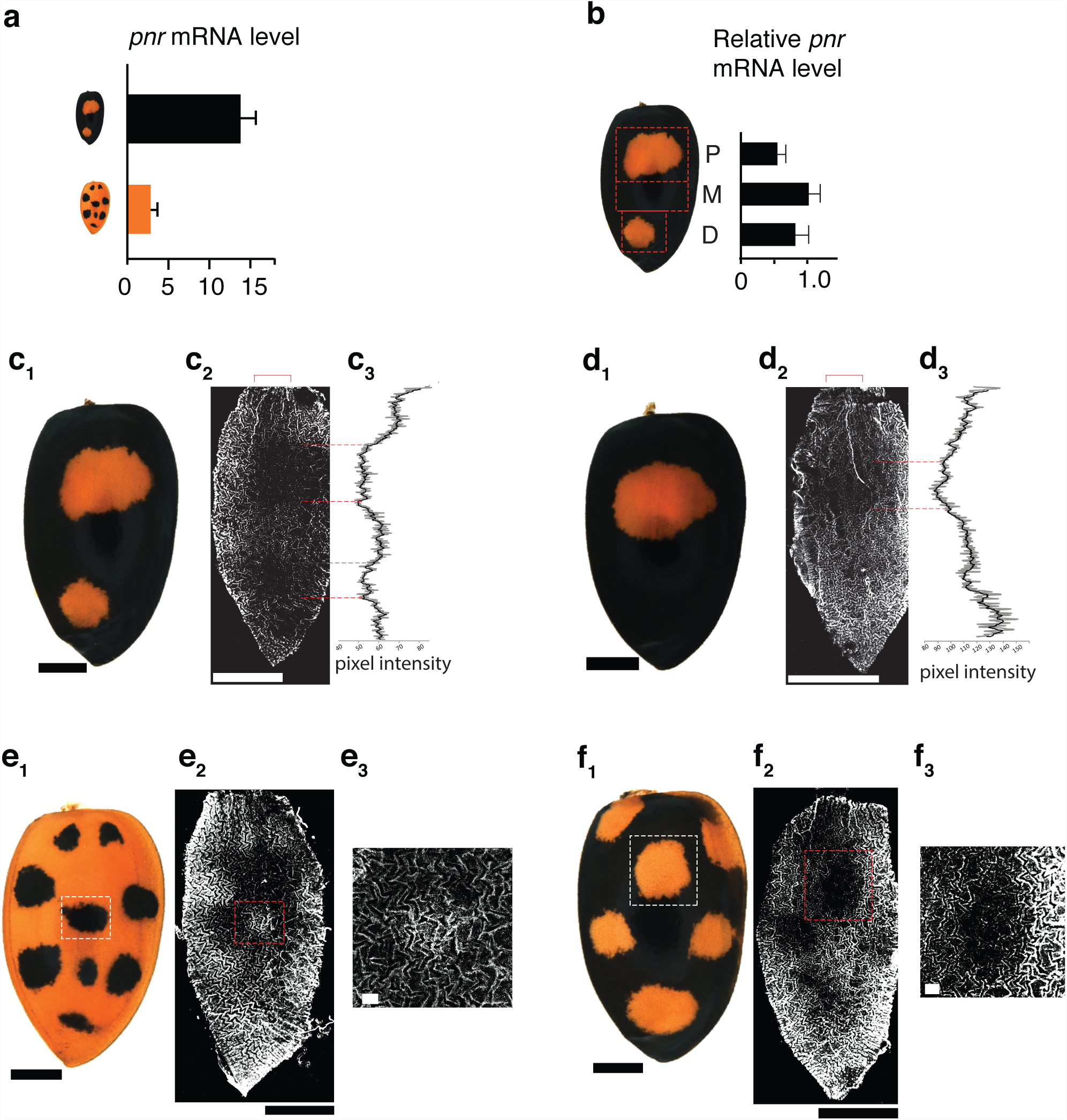
*pannier* expression pattern determines melanic colour pattern in each *H. axyridis* form.). **a**, Quantitative PCR of pannier mRNA from whole elytra reveals difference between Black-4Spots and Red-nSpots forms (two-tailed-t-test; n≥6; p=0.0005).). **b**, pannier is expressed at different levels in the presumptive red or black elytral areas in the Black-4Spots form (P, proximal, M, medial, D, distal), (n=5; two-tailed paired-t-test, p=0.02 (P vs. M), p=0.005 (M vs. D)). **c-f**, Immunodetection of Pannier protein in each colour form (~96h after pupation); (**c1-f1**) show adult elytron, (**c2-f2**) show anti-Pannier staining, (**c3, f3**) show pixel intensity along the PD axis of the elytron in the region delineated with a bracket in **c2** and **d2**), (**e3**, **f3**) show higher magnification of the selected areas (**e2**, f2). Note the reduction of signal intensity in the presumptive red/orange areas in each form. Scale bar; 1mm; 100µm for insets

In addition to allelic variation, colour pattern diversity in *H. axyridis* is shaped by the dominance relationships among colour form alleles^12^. Indeed, similarly to other species (e.g.,^16^), heterozygous individuals resulting from the cross of distinct homozygous *H. axyridis* forms produce black pigmentation in any part of the elytra that is black in either parental form^12^. Our results help to understand this phenomenon known as mosaic dominance at the molecular level. Since elytral Pannier expression patterns mimics adult black pigmentation patterns, and since each *pannier* allele carries its own *cis*-regulatory determinant to drive specific expression pattern, the expression pattern of *pannier* in heterozygotes will reflect the sum of individual form patterns. In other words, the mosaic expression of *pannier* during development, driven by the heterozygous alleles, produces a mosaic pigmentation pattern in adults (Fig. 4). This phenomenon compounds the effect of *pannier* allelic variation, increasing the complexity of colour patterns polymorphism in *H. axyridis*.

**Figure 4.**
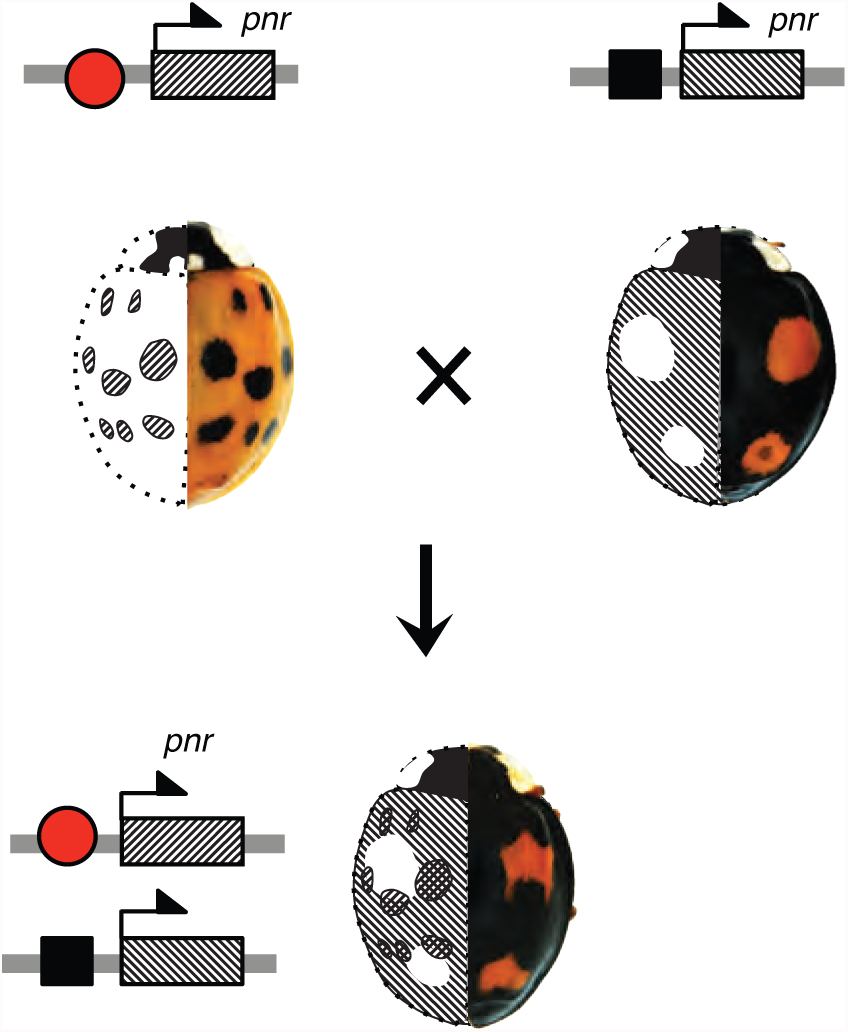
Genetic basis of the mosaic dominance¶. Each colour pattern form is determined by a particular cis-regulatory allele of *pannier,* symbolised with red square and black oval (top). In heterozygous individual (bottom), the mosaic colour pattern reflects *pannier* expression pattern, which is the sum of the two allelic forms. For each individual, the left elytron shows *pannier* expression pattern, and the right elytron shows the corresponding adult pigmentation pattern.

Finally, in order to precisely compare the sequences of *pannier* alleles between the Red-nSpots and another form, we sequenced the Black-4Spots form *de novo*. We chose the Black-4Spots form because it seemed the most divergent when compared to the Red-nSpots, based on genome-wide association results (Extended Data Fig. 1 and 2) and careful inspection of read coverage of the pools representative of colour forms (Supplementary Information). We generated the Black-4Spots draft assembly (*HaxB4)* from Illumina sequencing reads (Extended Data Table 1) and supplemented it by targeted BAC sequencing to derive a high quality 2.87 Mb sequence (% N = 0.96) spanning *pannier* and adjacent genomic regions. Strikingly, we found that the Red-nSpots and Black-4Spots sequences of the 5’ non-coding DNA and the first intron of *pannier* align poorly, in contrast to adjacent regions (Extended Data Fig. 7). Furthermore, we detected the footprint of a ca 50 kb long inversion within the first intron of pannier (Fig. 1d). This comparison indicates that the sequences of the non-coding DNA of *pannier* have diverged extensively between these forms.

Altogether, our results conclusively show that *pannier* plays a key role in the specification and the diversification of the main colour pattern forms in *H. axyridis*. *pannier* has never been reported to play a role in pigmentation in insects. It has therefore been co-opted for this function in the lineage leading to *H. axyridis*, presumably through the evolution of new regulatory connections with downstream effector genes directly involved in black pigment production^17^. This contrasts with other insect groups, including butterflies and fruitflies, in which different regulatory genes have been co-opted to generate wing colour patterns^3,6,17,18^. Furthermore, we showed that polymorphic colour patterns in *H. axyridis* arise from differential regulation of *pannier* spatial expression. The divergent non-coding sequences we have identified in the *H. axyridis* colour pattern forms are quite large (ca. 170 kb) and may host multiple discrete cis-regulatory elements. We propose that these *cis*-regulatory elements produce different *pannier* expression patterns and, ultimately, discrete melanic patterns, by interpreting differently a regulatory blueprint that is common to all *H. axyridis* colour pattern forms. This model is reminiscent of the mechanisms underlying thoracic bristles or wing pigmentation patterns diversity in *Drosophila* species^17,19,20^, and that has been proposed to explain wing colour pattern evolution in butterflies^6^.

We have demonstrated that the *cis*-regulatory region of *pannier* is highly divergent between the Red-nSpots and Black-4Spots alleles and that it includes a ca. 50 kb inversion. Furthermore, our data provides evidence of large-scale sequence variation among all four alleles of the main colour pattern forms in natural populations (Supplementary Information). We hypothesize that the numerous, rare colour pattern forms that have been described in *H. axyridis*^12^, are also determined by *pannier cis*-regulatory variation, and that they might result from rare mutational events, including rare recombination between the alleles of the main forms.

The striking sequence divergence among the *pannier* alleles of the main colour forms brings into question their evolutionary origin. Possibilities include ancient mutational events or even across species introgression events^10,21^. Thorough characterization of the *pannier* genomic region from different colour pattern forms both within *H. axyridis* and across coccinelid species (especially those harboring colour pattern polymorphisms^7^) will illuminate the evolutionary origin of the genomic determinants of colour pattern forms in *H. axyridis* and other ladybird species.

Finally, if sequence divergence likely helps preserving distinct *pannier* alleles by reducing recombination among them, selective mechanisms are also suspected to maintain different colour forms in natural populations of *H. axyridis* and affect their frequencies. Both local adaptation to climatic factors and seasonal variations in temperature have been suggested to affect colour forms proportions in space and time, possibly mediated by mate choice^10,11,22^.The identification of the genomic basis of colour pattern polymorphism will help to better characterize the evolutionary mechanisms that shape the striking colour pattern diversity in natural populations of *H. axyridis*, and to reveal potential pleitotropic effects of *pannier* alleles on traits involved in survival and reproduction^22,23^.

## Acknowledgments

We thank G. Beydon for constructing and screening the BAC library, L. Sauné and M. Galan for assistance in MISEQ sequencing, E. Lombaert, I. Goryacheva, Robert Koch, Jonathan Lundgren and C.E. Thomas for help in sampling wild *H. axyridis* populations, Ashraf Tayeh for assistance in lab-rearing of *H. axyridis* and Anthony Bretaudeau for making the *H. axyridis* genome assemblies publicly accessible. We are grateful to Nicolas Gompel, Nicolas Rode, Ruth Hufbauer and Renaud Vitalis for comments on the manuscript. A.E., B.F., J.F. and M.G. acknowledge financial support from the national funding agency ANR (France) through the European Union program ERA-Net BiodivERsA (project EXOTIC), and the GenoToul bioinformatics platform Toulouse Midi–Pyrenees for providing computing resources; B.P. and J.Y. acknowledge financial support from the CNRS, the European Research Council under the European Union’s Seventh Framework Programme (FP/2007-2013) / ERC Grant Agreement n° 615789, and the France-BioImaging infrastructure supported by the Agence Nationale de la Recherche (ANR-10-INSB-04-01, call “Investissements d’Avenir”), and the support of the NIG supercomputer at ROIS National Institute of Genetics (Japan); C.L.P., C.D. and M.M. acknowledge financial support from France Génomique National infrastructure, funded as part of “Investissement d’avenir” program managed by Agence Nationale pour la Recherche (contract ANR-10-INBS-09).

## Author contribution

M.G. conceived the project, designed the study, carried out bioinformatics and statistical treatments for genome-wide association studies and the identification of *HaxR* autosomal contigs, performed or supervised bioinformatic analyses and BAC contig construction, and wrote the manuscript; J.Y. carried out larval RNAi, cDNA sequencing, qPCR, immunohistochemistry and imaging studies, processed various bioinformatics treatments and wrote the manuscript; J.F. helped design the study and contributed to obtaining and maintaining various *H. axyridis* populations in the lab; A.L. processed molecular work for the *de novo* assembly, genome-wide association studies and BAC library PCR screening; A.A. contributed to obtaining and maintaining various *H. axyridis* populations in the lab; B.F. helped design the study and obtained funding; B.G. analysed the quality of the *de novo* genome assemblies and annotated the coding genes; J.L. designed the MinION study and processed associated bioinformatics treatments leading to the *HaxR de novo* assembly; E.L. processed bioinformatics treatments for the *HaxB4 de novo* assembly; H.P. and D.S. produced pool-seq and individual NGS data; C.L.R., C.D. and M.M. helped designed the MinION study, produced the MinION data and processed upstream bioinformatics treatments; H.B. developed and helped screening the Black-4Spots BAC library; K.G. helped design the study and produced some NGS data to construct the *HaxB4 de novo* assembly; L.L.H. provided *H. axyridis* individuals from Japan and contributed to drafting the manuscript; L. S.Z. provided *H. axyridis* individuals from China; H. V. helped design the study, obtained funding, produced RNA-seq data and genomic resources for the *HaxB4 de novo* assembly; B.P. and A.E. designed and directed the project, obtained funding, interpreted results and wrote the manuscript. All authors commented on the manuscript.

## Methods

### Description and naming of the four colour pattern forms that predominate in frequency in natural *H. axyridis* populations

(i) form *conspicua* (hereafter named “Black-2Spots” for clarity) has black background colour of elytra with two orange spots on each elytron and the top one larger than the bottom one. (ii) f. *spectabilis* (hereafter “Black-4Spots”) has black background of elytra with one large orange spot in the top-centre of each elytron, (iii) f. *axyridis* (hereafter “Black-nSpots”) has black background colour of elytra with many orange spots, and (iv) f. *succinea* (hereafter “Red-nSpots”) has orange to red background colour of elytra with the number of black spots ranging from 0 to 19. See Fig. 1a of main text for illustrations.

### De novo assembly of the *Harmonia axyridis* genome from individuals of the colour pattern form Red-nSpots

Four Oxford Nanopore Technologies (ONT) libraries were prepared using the Ligation Sequencing Kit 1D (SQK-LSK108 ONT), according to the manufacturer’s protocol 1D Genomic DNA by ligation (SQK-LSK108). Briefly, 60 µg of genomic DNA was extracted using the Genomic-tip 500/G kit (Qiagen) from a pool of thorax tissue belonging to 12 Red-nSpots males from a lab-reared population founded by Red-nSpots individuals originating from the biocontrol population BIOTOP (France). The DNA sample was divided into four aliquots and sheared into 25 kb (n=3) or 30 kb (n=1) fragments using the Megaruptor system (Diagenode). One of the 25 kb sheared DNA sample was additionally size-selected prior to library preparation using the BluePippin system (Sage Science, Beverly, USA) to remove fragments smaller than 10 kb. A DNA Repair (NEBNext FFPE Repair Mix M6630) step as well as an End-repair and dA-tail step (NEBNext End repair / dA-tailing Module E7546) were then processed on 0.34 pmols of the sheared DNA sample, followed by ligation of sequencing adapters. Then 0.07 pmols of library were loaded onto an R9.5 flow cell containing at least 800 active pores and run for 48 hrs, on a MinION sequencer (ONT) and sequenced by the Minknow ONT software (v1.7.10 or v1.7.14). Base calling was carried out using Albacore (from ONT, v1.2.4 or 2.0.2) with default settings yielding 2.46×10^6^ reads (22.9 Gb) (Supplementary Table 1) that were trimmed using Porechop 0.2.3^1^ with default options. Trimmed reads (Nanofilt v. 1.1.3,^2^) with a quality score Q>9 and longer than 500 bp (1.34×10^6^ reads; 17 Gb) were combined and further self-corrected using canu v.1.6^3^ with default options. Genome assembly was then performed based on the corrected long reads using SMARTdenovo v.1.0.0 run with default settings. A last polishing step with Pilon v1.22^4^ was carried out using paired-end (PE) Illumina sequencing reads (ca. 100X coverage) available for two pools of Red-nSpots individuals (see below). The resulting assembly, denoted *HaxR*, consisted of 1,071 contigs spanning 429 Mb (N50 = 1,434 Mb) encompassing 97.2% of the BUSCO (Benchmarking Universal Single-Copy Orthologs^5^) highly conserved arthropod gene set. See Extended Data Table 1 for additional *HaxR* statistics. The *HaxR* assembly is publicly available at http://bipaa.genouest.org/sp/harmonia_axyridis.

### Identification of *HaxR* autosomal contigs using a female to male read mapping coverage ratio

Barcoded DNA PE libraries with an insert size of ca. 550 bp were prepared using the Illumina TruSeq Nano DNA Library Preparation Kit following the manufacturer’s protocols using six DNA samples extracted from three Red-nSpots males and three Red-nSpots females originating from a wild *H. axyridis* population from Mississippi (USA). Libraries were then validated on a DNA1000 chip on a Bioanalyzer (Agilent) to determine size and quantified by qPCR using the Kapa library quantification kit to determine concentration. The cluster generation process was performed on cBot (Illumina Inc.) using the Paired-End Clustering kit (Illumina Inc.). Each individual library was further paired-end sequenced on a HiSeq 2500 or 2000 (Illumina, Inc.) using the Sequence by Synthesis technique (providing 2×125 or 2×100 bp reads, respectively) with base calling achieved by the RTA software (Illumina Inc.). After removal of sequencing adapters, reads were mapped onto the *HaxR* assembly using default options of the mem program from the bwa 0.7.12 software package^6^. Read alignments with a mapping quality Phred-score <20 and PCR duplicates were further removed using the *view* (option *-q 20*) and rmdup programs from the samtools 1.3.1 software^7^, respectively. Read coverage at each contig position for each individual sequences was then computed jointly using the default options of the samtools 1.3.1 depth program. To limit redundancy, only one count every 100 successive positions was retained for further analysis and highly covered positions (>99.9^th^ percentile of individual coverage) were discarded. The estimated individual median overall coverage ranged from 6 to 21 (see Supplementary Table 2 for details).

To identify autosomal contigs, we used the ratio *ρ* of the relative (average) read coverage of contigs between all females and all males (weighted by the corresponding overall genome coverage) expected to equal 1 for autosomal contigs and 2 for X-linked contigs^8^. Contigs smaller than 100 kb were discarded from further analyses because they displayed a high coverage dispersion of their coverages (Supplementary Table 3) together with 12 of the remaining contigs with extreme values of *ρ* (*ρ* < 05 or *ρ* > 2.5). We further fitted the *ρ* distribution of the 492 remaining contigs (398 Mb in total) as a Gaussian mixture model with two classes of unknown means and the same unknown variance. The latter parameters were estimated using an Expectation-Maximization algorithm as implemented in the mixtools R package^9^. As expected the estimated mean of the two classes µ_1_=0.96 and µ_2_=1.90 were only slightly lower (see Supplementary Table 3 for further details) to that expected for autosomal and X-linked sequences allowing to classify the different contigs. We could hence therefore assign classify with a high confidence (p-value < 0.01) 457 contigs as autosomal (377.5 Mb in total) and 18 contigs as X-linked (16.85 Mb in total) with high confidence.

### Genome-wide scan for association with the proportion of Red-nSpots individuals using Pool-Seq data on 14 population samples

Barcoded DNA PE libraries with insert size of ca. 450 bp were prepared using either the Illumina Truseq DNA sample prep kit (n=2) or the Nextera DNA Library Preparation kit (n=12) following manufacturer protocols using 14 DNA pools (each pool including the head - or the leg for some pools - from n = 40 to n = 100 individuals) collected in eight populations representative of the world-wide genetic diversity^10^ and the four main colour pattern forms of the species (Extended Data Table 2). Illumina sequencing, processing and mapping of reads to the *HaxR* assembly was performed as described above for individual data (see Supplementary Table 4 for further details). The 14 Pool-Seq BAM files were processed using the mpileup program from the samtools v1.3.1 software^7^ with default options and -*d 5000* and *-q 20*. Variant calling was then performed on the resulting mpileup file using VarScan mpileup2cns v2.3.4^11^ (options --min-coverage 50 --min-avg-qual 20 --min-var-freq 0.001 --variants -- output-vcf 1). The resulting VCF file was processed with the vcf2pooldata function from the R package poolfstats v0.1^12^ retaining only bi-allelic SNPs covered by > 4 reads, < 99.9^th^ overall coverage percentile in each pool and with an overall MAF > 0.01 (computed from read counts). In total, 18,425,210 SNPs mapping to the 457 autosomal contigs were used for genome-wide association analysis with a median coverage ranging from 18 to 45X per pool (Supplementary Table 4).

Genome-wide scans for association with the proportion of individuals of a given colour pattern form in each pool were performed using the program BayPass 2.1^13^. Capitalizing on the large number of available SNPs, we sub-sampled by taking one SNP every 200 SNPs along the genome, dividing the full dataset into 200 sub-datasets (each one including ca. 92,500 SNPs). These sub-datasets were further analysed in parallel under the BayPass core model using default options for the Markov Chain Monte Carlo (MCMC) algorithm (except -npilot 15 -pilotlength 500 -burnin 2500). Three independent runs (using the option -seed) were performed for each dataset. The estimated model hyper-parameters were highly consistent across both runs and datasets. Support for association of each SNP with the corresponding prevalence covariate was then evaluated using the median Bayes Factor (BF) computed over the three independent runs. BFs were reported in deciban units (db) with 20 db corresponding to 100:1 odds, 30 db to 1000:1 odds, and so on^14^.

### *de novo* sequencing of the Black-4Spots colour pattern allele

Starting from Black-4Spots individuals from the low diversity BIOTOP biocontrol population (France), we carried out four generations of full-sib crossings to produce a Black-4Spots inbred line, hereafter referred to as B4sIL. This aimed to improving further assembly steps by reducing the overall genetic variability. Four DNA PE libraries with insert sizes of ca. 250 bp (n=2), 400 bp (n=1) and 600 bp (n=1) and two DNA Mate Pairs (MP) libraries with insert sizes of ca. 3 kb and 8 kb were constructed from B4sIL DNA (3-4 individuals per library) using standard Illumina (Inc.) kits; and 2 Long Jumping Distance libraries (Eurofins MWG Operon) with insert sizes of ca. 3 kb and 8 kb. All these libraries were sequenced on a HiSeq2500 sequencer with either 2×100 bp or 2×150 bp reads (Supplementary Table 6). Raw reads were filtered for bacterial and human sequence contaminants using deconseq^15^ and trimmed using Trimmomatic v0.22^16^. Genome assembly was then performed using AllPath-LG^17^ with default options except Haploidify=True to account for residual polymorphism in the sequenced individuals. This led to a first assembly consisting of 6,883 scaffolds (N50 = 921 kb) totaling 378 Mb (%N = 14.8). To further improve the assembly, we generated long reads using the Pacific Biosciences RS II platform. To that end seven SMRT bell libraries were prepared using size fractionated (shearing size of 25 kb and size selection cut-off of 10 kb) high molecular weight DNA prepared from B4sIL individuals and loaded into SMRT Cell 8Pac v3 for sequencing on Pacific Biosciences RS II system with P6C4 chemistry by Treecode (Malaysia). The seven resulting sequence movie files were processed and analysed using Pacific Biosciences SMRT Analysis Server v2.3.0 with default settings. After the filtering step, a total of 422,222 reads (N50 = 21,521 kb) were used to carry out gap filling and scaffolding using PBJelly v1.3.1^18^. The final Black-4Spots assembly, referred to as *HaxB4*, consisted of 6,586 scaffolds (N50=978.4 kb) totaling 393 Mb (%N=5.84). Both genome-wide association studies conducted as described above but with *HaxB4* as a reference (Supplementary Fig. 1 and 2) and sequence alignment of the utg686 contig of the *HaxR* assembly using Mummer^19^ (and *BLAST+*^20^) tools allowed unambiguous identification of a 5.96 Mb HaxB4 scaffold that included the colour pattern locus and adjacent genomic regions.

Because the *HaxB4* assembly remained less contiguous and accurate than the *HaxR* assembly (see Extended Data Table 1 for a comparison), we performed a finishing step relying on a newly developed BAC (Bacterial Artificial Chromosomes) physical map covering the Black-4Spots allele of the colour pattern locus. To that end, a BAC library of 13,824 BACs (141+/-41 kb insert size after sizing of 48 BACs) with ca. five genome equivalent coverage was constructed in the vector pIndigoBAC-5 using high molecular weight DNA extracted from 300 B4sIL larvae (of developmental stage L1), as described in^21^. The BAC library, deposited at the CNRGV (INRA, Toulouse, France. https://cnrgv.toulouse.inra.fr/Library/Asian-ladybird), was organized in 2-dimension pools for PCR screening. A total of 98 PCR primers pairs designed against the *HaxB4* assembly or newly generated BAC-end sequences were screened on the library. This allowed defining a Minimum Tiling Path of 14 BACs covering 1.9 Mb for local correction of scaffold mis-assembly (Supplementary Fig. 2). Shotgun sequencing was carried out using the Pacific Biosciences RS II system and Illumina MiSeq sequencer for 3 and 10 BACs, respectively^21^. Finally, we manually edited the *HaxB4* assembly using the BAC sequences to derive a high quality 2.87 Mb sequence (% N = 0.96) including the candidate region of the Black-4Spots allele of the colour pattern locus as well as adjacent regions (Supplementary Data 1). Alignment of the later sequence to the *HaxR* assembly (i.e. Red-nSpot allele) using nucmer from the package Mummer^19^ allowed the scaffolding of the five neighboring *HaxR* contigs (Supplementary Fig. 4) represented in Fig. 1c.

### Larval RNAi

We synthesized double-stranded RNAs (dsRNA) with T7 polymerase as described previously^22^. DNA fragments for the transcription were amplified by PCR using primers containing T7 polymerase promoter sequences at their 5’ends (see Supp Data Table 6 for primer sequences). We used cDNA from Black-4Spots or Red-nSpots forms. Sense and antisense transcripts were simultaneously synthesized using RiboMax express RNAi System (Promega), annealed, treated with RQ1 DNase (Promega), and precipitated with ethanol. The quality of dsRNA was examined by agarose gel electrophoresis, and the concentration was roughly measured by spectrophotometer ND-1000 (NanoDrop Technologies), and 2 µg/ul in nuclease free water were used for injection. Larvae were anesthetized just before pupation on a CO_2_ pad, and 400-600 nl of dsRNA was injected into hemolymph using Nanoject (Drummond Scientific).

### cDNA sequencing

Fragments of *pannier* for Black-4Spots and Red-nSpots forms were amplified separately by PCR from cDNA of elytra of Black-4Spots or in Red-nSpots forms. Total RNA was extracted using TRI reagent (Invitrogen), followed by DNase I treatment. cDNA was generated using First Strand cDNA Synthesis Kit (New England BioLabs). Resulting PCR fragments were approximately 1.8 kbp in length for both melanic forms when using the primer set (Ha_pnrF1; CGGTACGAGATAAGCGAATAAGG, Ha_pnr-F1; TTACCATTTACAAATATATTTACATGGTTGTTG). Each PCR product was inserted into cloning vector pGEM-T Easy (Promega) for Sanger sequencing.

### Quantitative PCR (qPCR)

Total RNA was extracted from whole elytron of homozygous Black-4Spots (n=7) or Red-nSpots (n=6), or dissected elytron of homozygous Black-4Spots (n=5) at late pupal stage (96 hr after pupation) with TRI reagent (Invitrogen). RNA samples were reverse transcribed using First Strand cDNA Synthesis Kit (New England BioLabs). We omitted DNase I treatment, because all pairs of forward and reverse primer for qPCR were designed in different exons of each gene, which are separated by long introns. Furthermore, for accurate comparison, we confirmed the absence of nucleotide substitutions in the primer sequences in the different colour pattern forms. qPCR and data analysis were performed on StepOne Real-Time PCR System (Applied Biosystems) with Power SYBR Green Master Mix (Applied Biosystems). The data was normalized using eukaryotic initiation factor 4A (eIF4A) and 5A (eIF5A), and statistical significance of expression differences was established using two-tailed-*t*-test. All primer sets and each R^2^ value of standard curves are listed in Supp. Table 6.

### Immunohistochemistry

An antibody against *H. axyridis* Pannier (Ha_Pannier) was raised by Genscript, using as an antigen the first 384 amino acids of the protein. To test that this antibody recognizes Ha_Pannier we ectopically expressed Ha_Pannier using the Gal4/UAS system in *Drosophila melanogaster*. Specifically, we stained *engrailed*-Gal4, UAS-GFP; UAS-Ha_Pannier larval imaginal disks with anti-Ha_Pannier and anti-GFP using standard procedures. We observed co-localization of the signal in the posterior compartement of the disk, as expected, showing that the Ha_Pannier antibody recognizes Ha_Pannier *in vivo*.

Late pupal *H. axyridis* elytra are covered with a cuticle layer that is impenetrable for antibodies. Therefore, before staining, we split each elytron into two halves, separating the dorsal and ventral halves. For this we followed the protocol that has been developed for *Drosophila* wings^23^ with some modifications. Elytra were dissected from pupae at late stage (around 96 hr after pupation) in PBS, and fixed in 4% paraformaldehyde (5-10min at room temperature). The edges of the elytra were trimmed off with a razor blade before transferring the elytra on a piece of adhesive tape (Tesa). Another piece of adhesive tape was positioned on top of the immobilized elytra, and then gently removed to separate the two faces of the elytra. The two pieces of tape with split elytra (one is dorsal, the other is ventral side of the elytra) were fixed 4% paraformaldehyde again (1-5min at room temperature) and stained (overnight at 4°C) with anti-Ha_Pannier antibody at 1:70 dilution in 1% bovine serum albumin (BSA), followed by visualization with Alexa-dye-conjugated secondary antibodies (Invitrogen) at 1:100 dilution in 1% BSA (1h at room temperature). Cell nuclei were stained with DAPI. The pieces of tapes with stained elytra were mounted on microscope slides with VECTASHIELD (VECTOR).

### Imaging

Anti-Pannier stainings were imaged under LSM510 confocal microscope (Zeiss) with identical settings (e.g., objective lens, pinhole size, laser power, number of stacks, etc) for all samples. All raw confocal images were processed identically in ImageJ 1.51, and then enhanced separately with (Adobe Photoshop). Mean intensities of anti-Pannier signal were measured in rectangular sections using the Plot Profile command of ImageJ 1.51.

Adult *H. axyridis* (2 days post eclosion) were imaged on a Leica Z6Apo macroscope equipped with a ProgRes C5 ccd camera (Jenoptik). Several images were taken at different Z positions, and stacked together using HeliconFocus. Images were enhanced using Adobe Photoshop.

### Genomic sequence divergence and gene structure at the colour pattern locus

Genomic sequences of Red-nSpots (utg676) and Black-4Spots (HaxB4) contigs including *pannier* were visualized with dot plot using GenomeMatcher^24^. Conserved sequence blocks were further detected and visualized with Artemis Comparison Tool (ATC)^25^ for figure 1d. To identify reliable orthologous positions between the two contigs we first extracted long homologous blocks using blast2seq under high stringency parameters (blastn, e-value<0.01, alignment length≥2 kbp), and plotted them on the dot plot. The linear approximation was y=x+357764 (R^2^=0.973, y-axis; *utg676*, x-axis; *HaxB4*). There was blank region (where there are no plots) in the middle of this linear approximation, which corresponds to a highly diverged region (202 kb and 234 kb in *utg676* and *HaxB4*, respectively). We subsequently checked shorter homology blocks (≥1000 bp) around the breakpoints toward the center of the blank region to determine the borders more precisely. We considered continuous two or more conserved blocks (≥1000 bp) within 20 x 20 kb sliding window along the line approximation. Thus, we defined two breakpoints for the boundary between continuous conservation and divergent regions, the latter one spanning 173,272 bp on *utg676* (554,399 - 727,671), in line with the ca. 170 kb region delimited by our genome-wide association study, and 209,085 bp on *HaxB4* (913,592 - 1,112,677).

To identify gene structures around the divergent genomic region (173 kb on utg676), we mapped RNA-seq reads PRJEB13023^26^ (100bp paired-end, adult and larva *Harmonia* transcripts) deposited in Sequence Read Archive (SRA) to repeat-masked genome contigs using Tophat 2.1.0^27^, followed by assembling transcripts by Cufflinks v2.2.1^28^ using default parameters on the NIG supercomputer at ROIS National Institute of Genetics. Resulting genes were named after sequence homology to protein database of *Drosophila marlanogaster* (dmel-all-translation-r6.07) and *Tribolium_castaneum* (GCF_000002335.3_Tcas5.2). For the Black-4Spots sequence, since the number of mapped reads on exon-1 of *pannier* isoform-1 was low, we validated this gene structure using additional RNA-seq reads (Heiko V., unpublished). The mapped reads were further confirmed by eye on Integrative Genomics Viewer (IGV)^29^ to determine the gene structures.

---

**Extended Data Fig. 1:**
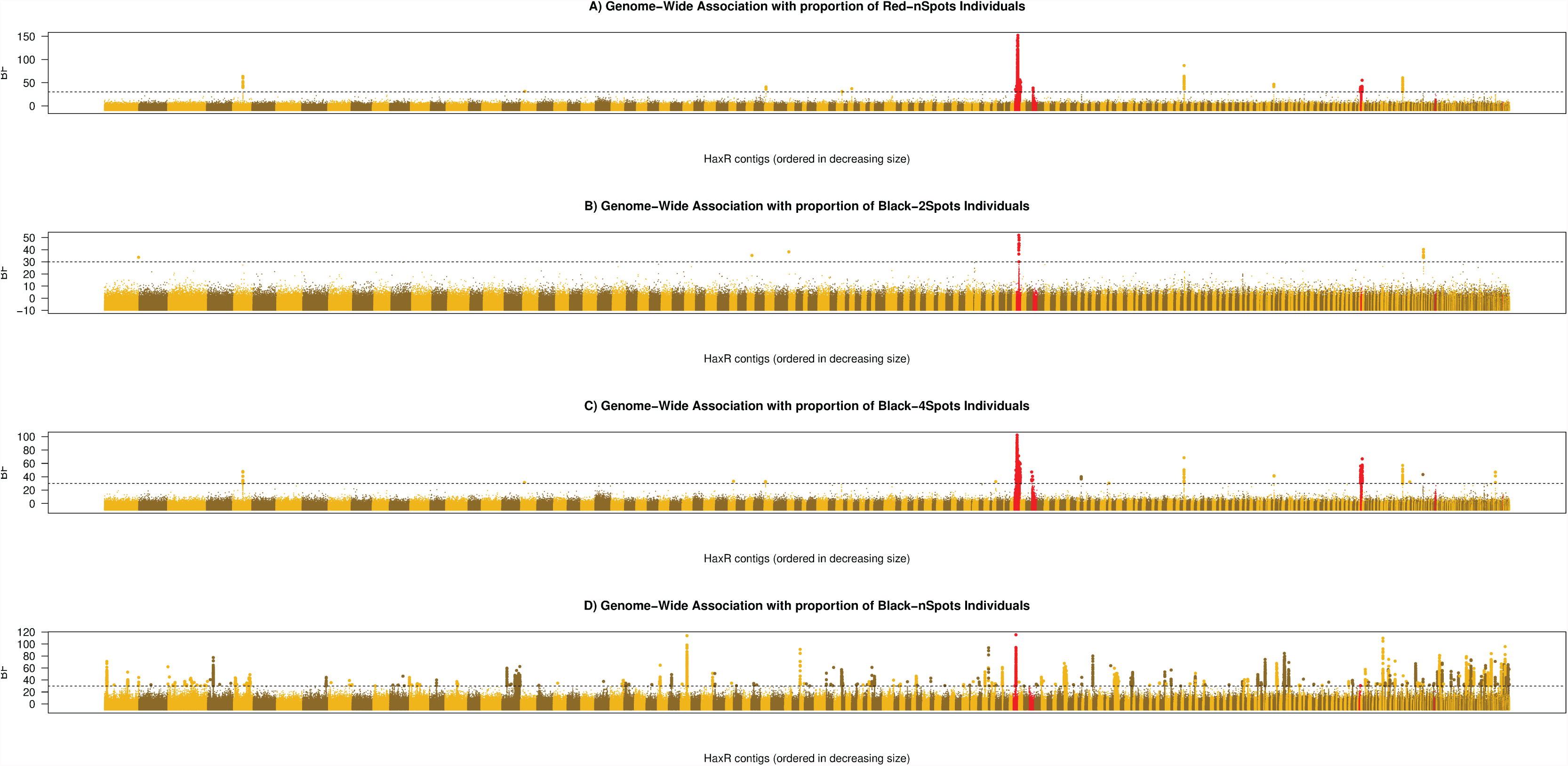
Results of the genome-wide association study using the assembly *HaxR* considering each of the four main colour pattern forms as covariate separately.

**Extended Data Fig. 2:**
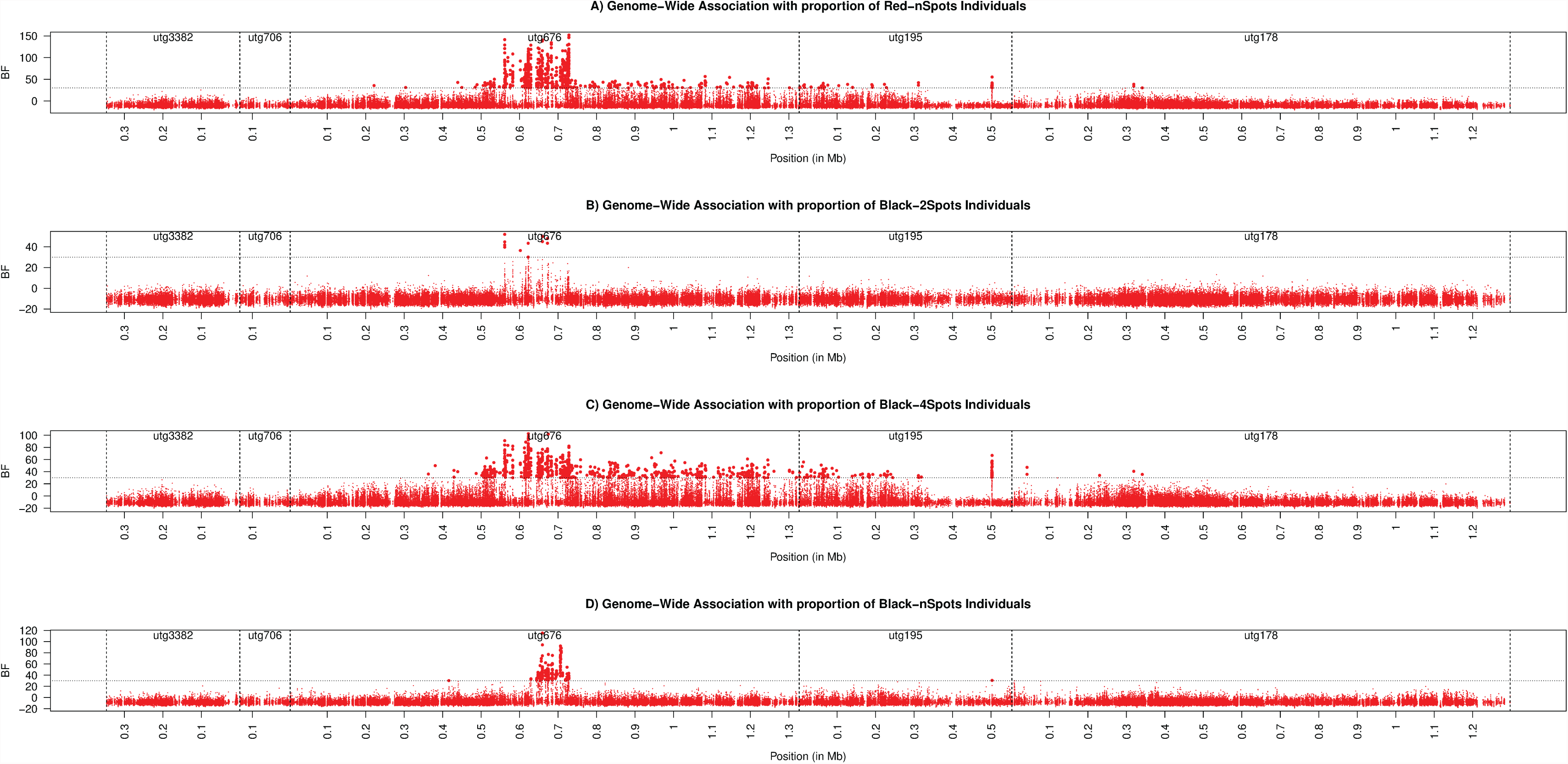
Results of the genome-wide association study focusing on the colour pattern genomic region of the assembly *HaxR* considering each of the four main colour pattern forms as covariate separately.

**Extended Data Fig. 3.**
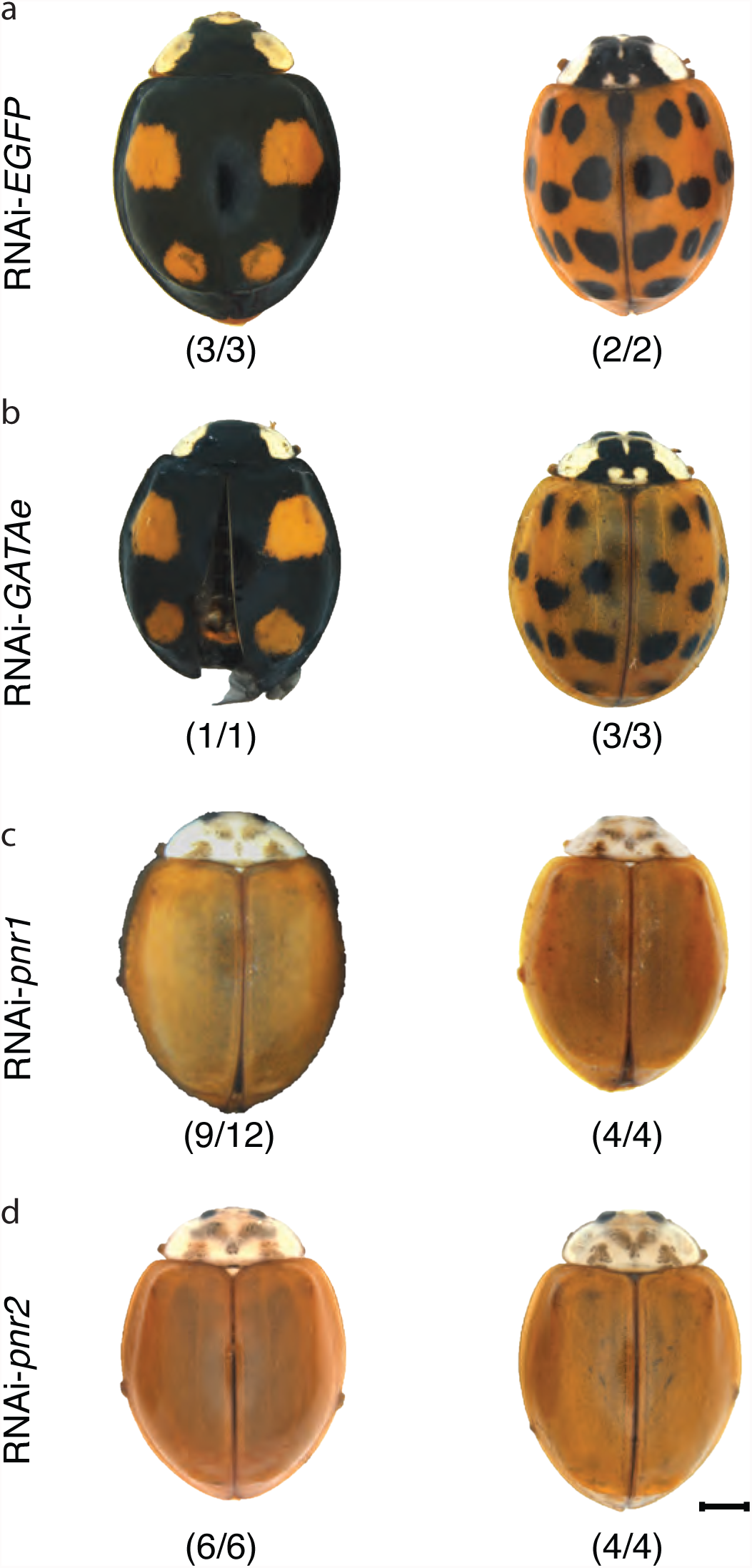
*pannier*, but not *GATAe*, is necessary for adult pigmentation patterns. RNAi phenotypes after larval injection of dsRNA targeting *eGFP* (negative control) (a), *GATAe* (b) or *pannier* (c, d) in Black-4Spots (left columns) or Red-nSpots (right columns) forms. *pnr1* and *pnr2* target different, non-overlapping regions of the *pannier* coding cDNA (shown in Extended Data Fig. 5). Knock down of *GATAe* had a strong effect on survival, with only few injected individuals producing viable adults. Numbers in parentheses indicate the proportion of eclosed adults with the representative pigmentation defect for each melanic form. Scale bar; 1mm

**Extended Data Fig. 4.**
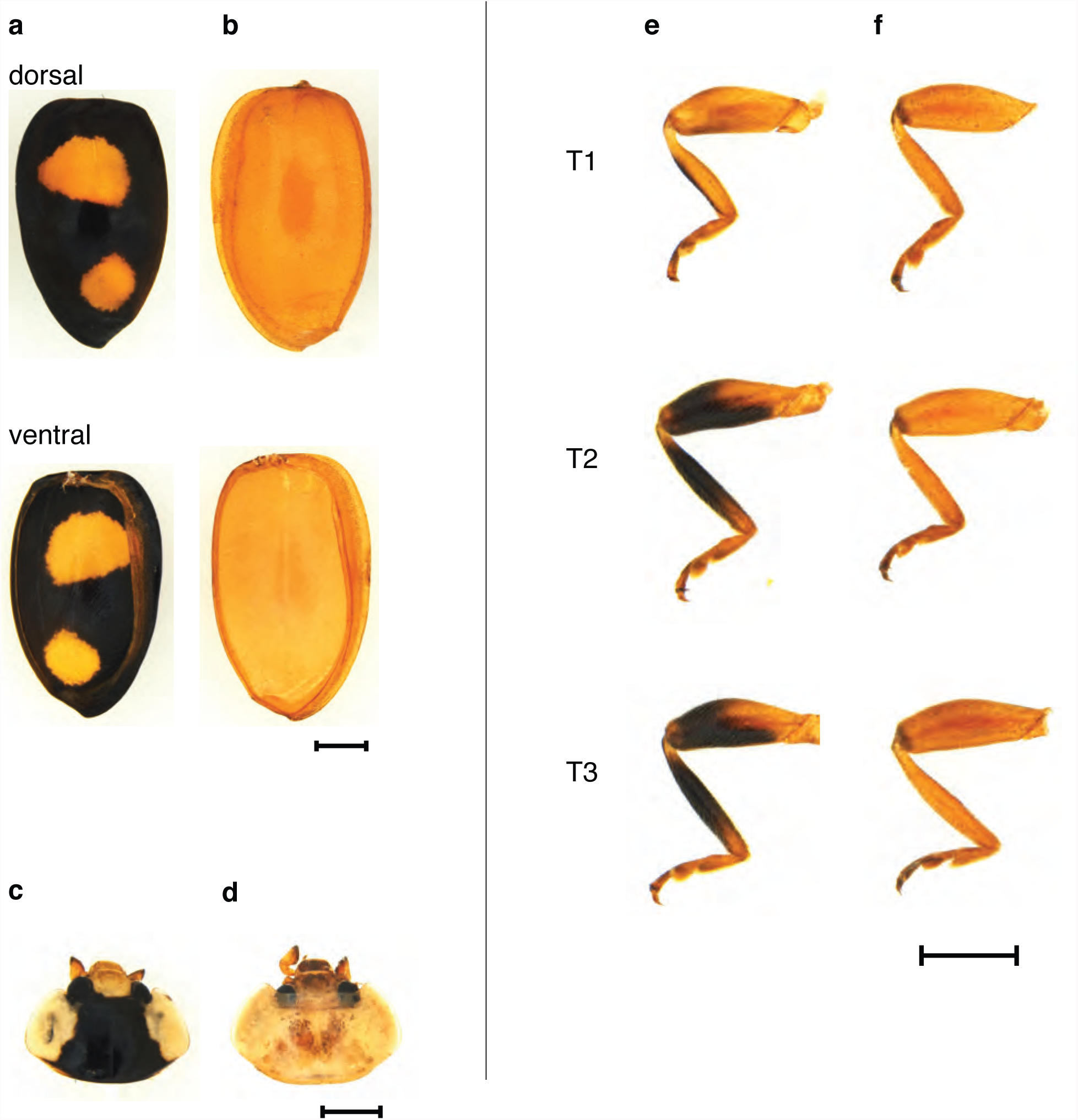
Details of RNAi-*pannier* phenotypes. Detailed pigmentation phenotypes between wild type and RNAi-pannier (right) in male Black-4Spots. Dorsal and ventral views of wild type elytra (a), or RNAi-*pannier*; (b). Head of wild type (c), or RNAi-*pannier* (d); T1, T2, T3 legs of wild type (e), or RNAi-*pannier* (f). Scale bars; 1mm.

**Extended Data Fig. 5.**
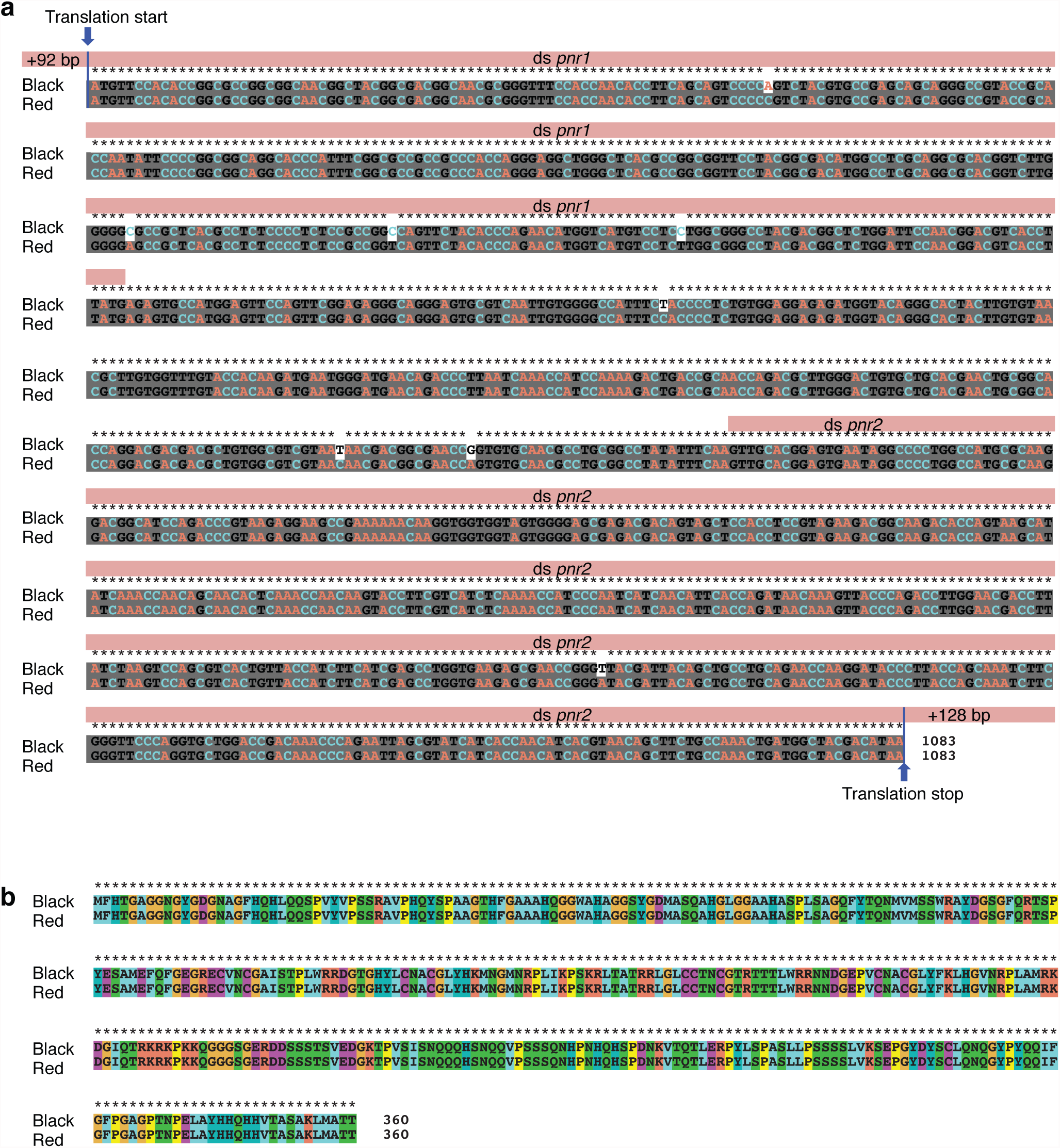
Alignment of *pannier* coding sequence from Black-4Spots and Red-nSpots forms. a, Nucleotide alignment of *pannier* coding sequences from two forms. Grey background indicates conserved sequences. The two DNA fragments amplified for dsRNAs synthesis are shown in pink. b, Protein sequences alignment. Note the absence of non-synonymous substitutions between Black-4Spots and Red-nSpots sequences.

**Extended Data Fig. 6.**
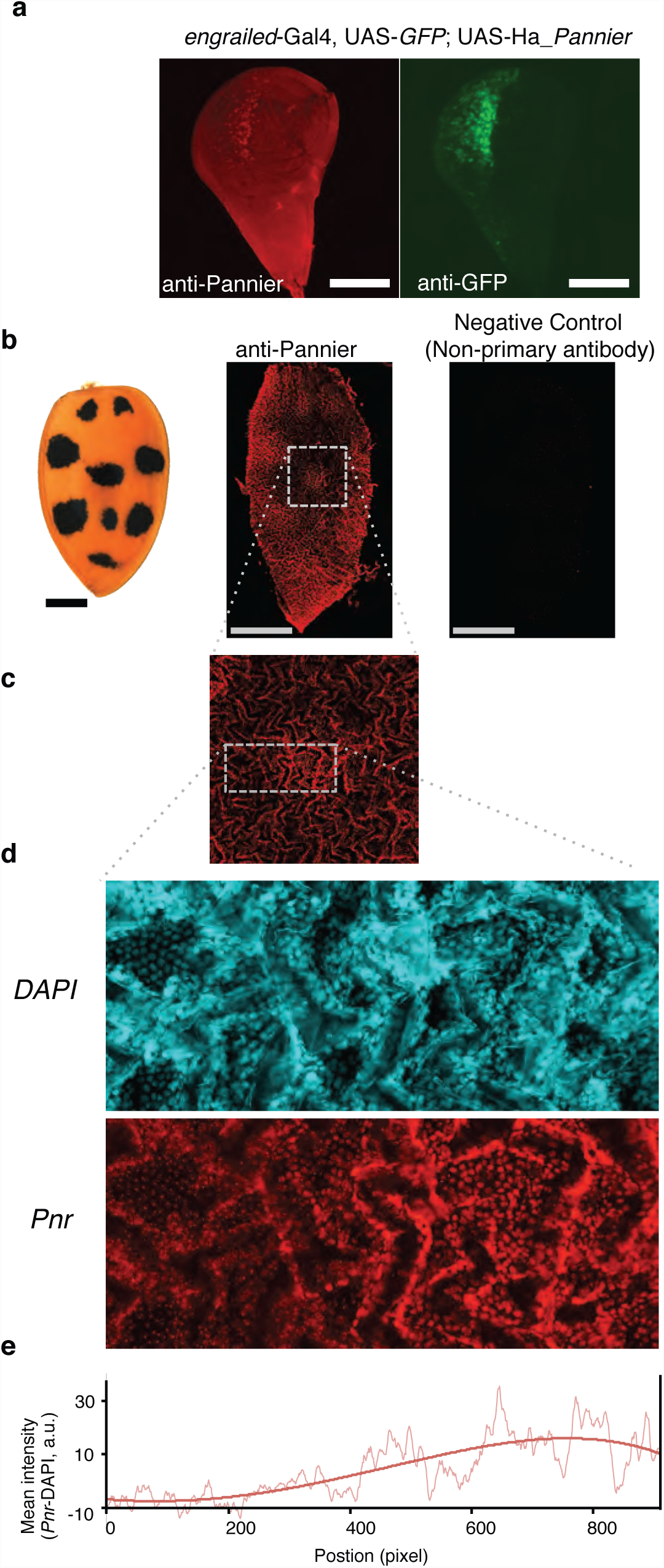
Details of immunohistochemistry with anti-pannier antibody. **a**, anti-Ha_Pannier and anti-GFP double staining on wing imaginal disk of *D. melanogaster engrailed-*Gal4, UAS-GFP; UAS-Ha_Pannier. The signals overlap in the posterior compartment of the disk, showing that the anti-Ha_Pannier recognizes Pannier protein in vivo. **b**, *H. axyridis* Red-nSpots elytron (left panel) and fluorescent image of immunohistochemistry with anti-Ha_Pannier antibody (central panel), or secondary antibody only (right panel). **c**, Enlargement of the region boxed in (**b**). d, Enlargement of the region boxed in (**c**) with DAPI staining. Scale bars, a: 100µm; b:1 mm, b,c; 100µm. Note that “jaggy lines” are observed on both DAPI and Ha_Pannier images, which may be due to folded structure of pupal elytra. **e**, Normalized mean intensities (intensity in anti-pannier staining (lower) - intensity in DAPI staining (upper)) at each position along x-axis and its polynomial fitting curve. In order to compensate for the tissue heterogeneity, we subtracted for each pixel the DAPI staining intensity from the anti-Pannier staining intensity.

**Extended Data Fig. 7.**
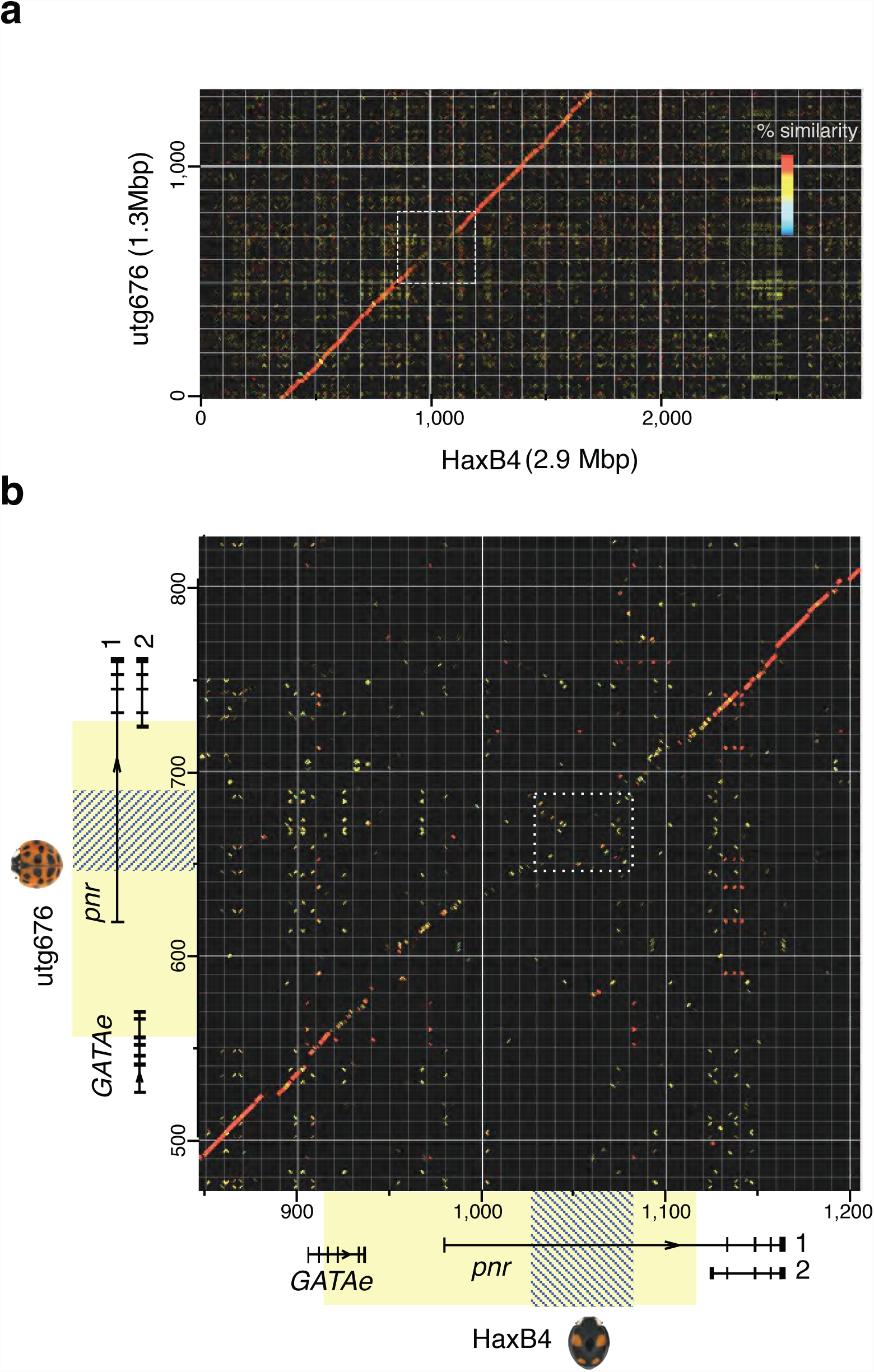
Comparison of utg676 and HaxB4 sequences **a**, Dot-plot between genomic scaffolds from the Black-4Spots (HaxB4) and Red-nSpots (utg676) forms. The most divergent sequence (white dashed box in a) in shown at a higher magnification in **b**). **b**, Traces of a sequence inversion are visible in the divergent region (white dashed box).

**Extended Data Table 1:** Statistics characterizing the assembly *HaxR* obtained from Red-nSpots individuals and the assembly *HaxB4* obtained from Black-4Spots individuals.

|  | Assembly *HaxR* | Assembly *HaxB4* |
| --- | --- | --- |
| Data | MinION long reads (65X)<br>Illumina PE reads (100X) | Illumina PE reads (65X)<br>Illumina MP reads (24X) |
| Assembler | SMARTdenovo | ALLPATH-LG |
| Nb. of sequences | 1,071 contigs | 6,586 scaffolds |
| Total length (Mbp) | 429 | 393.1 |
| Average length (Kbp) | 400.9 | 59.7 |
| Max size (Kbp) | 7,499 | 5,635 |
| Total Ns (bp) | 22 | 22,814,986 |
| N50 (Kbp) | 1,434 | 978.4 |
| BUSCO (complete) | 97.2 % | 86.0 % |
| BUSCO (fragmented) | 1.3 % | 8.7 % |
| BUSCO (missing) | 1.5 % | 5.3 % |

**Extended Data Table 2.**
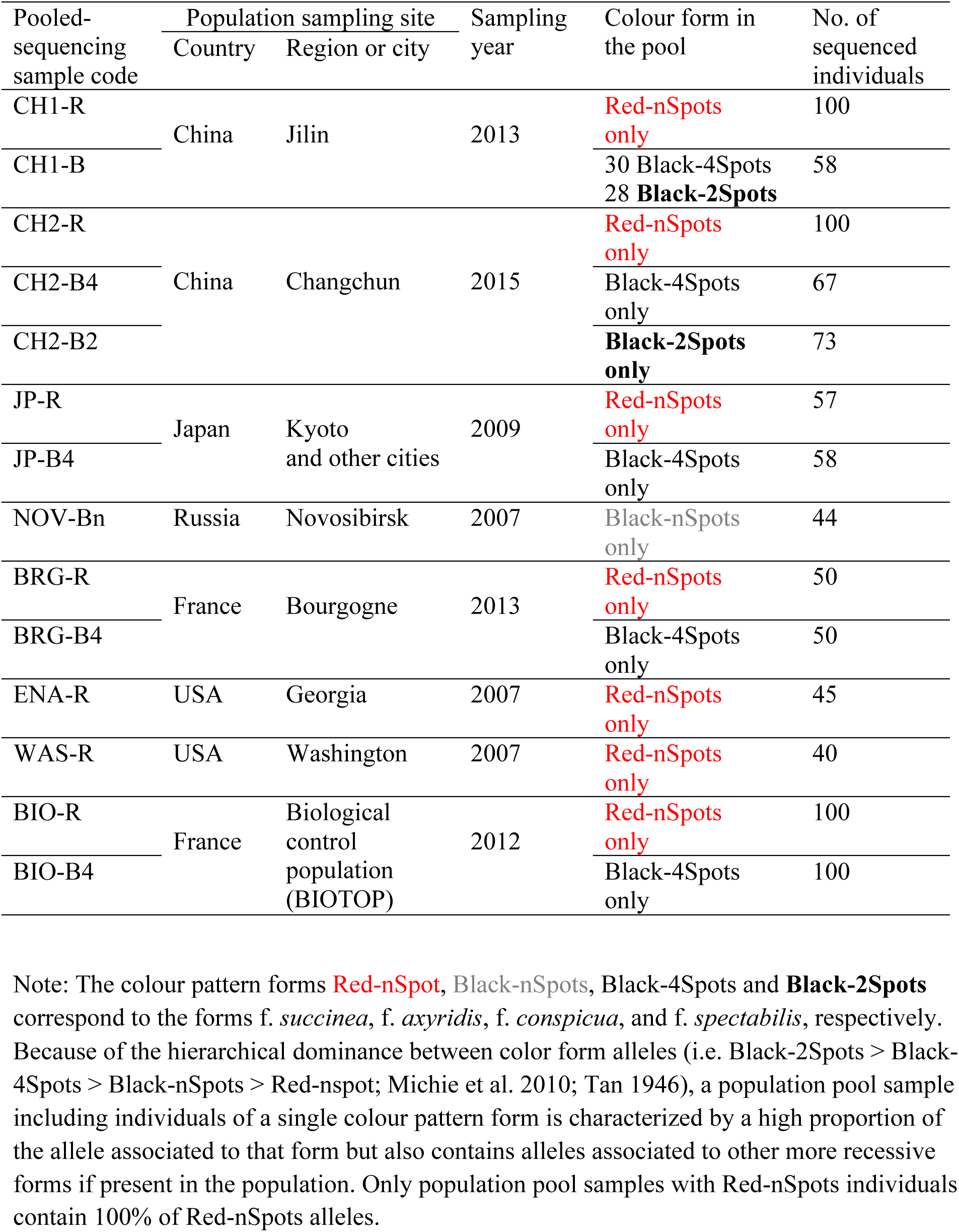
Sequenced pools of individuals representative of the world-wide genetic diversity and the four main colour pattern forms of *H. axyridis*

